# Predation at a snail’s pace. What is needed for a successful hunt?

**DOI:** 10.1101/420042

**Authors:** Wouterus T. M. Gruijters

**Affiliations:** Auckland Grammar School, 55 Mountain Road, Grafton, Auckland 1023, New Zealand

**Keywords:** Paryphanta busbyi, Gastropoda Rhytidae, Kauri snail, pupurangi, predator, humidity, wind, earthworm, tracking, lair, migration

## Abstract

Daily observations of wild *Paryphanta busbyi* (New Zealand Kauri snail) along the edge of a driveway were carried out for just over one year. Documented behaviours include the snail’s speed of movement, thrusting their heads down into leaf litter to wait in ambush for prey, the daily retreat into lairs, burrowing, and the grooming of the snail’s shell. The most significant environmental factors affecting the presence of the snails were humidity and wind direction. There was a migration of snails into and out of the observable study area at different times of year. Due to the snail’s pace of 1.9 to 3 m/hr relative to its prey at 5-10m/hr the snails would need to migrate to and inhabit areas where there is likely to be trackable prey. Overall the observations indicate the hunting strategy of a periodically active ambush predator sometimes based in a lair and migrating to suitably humid and windless areas as needed to allow tracking of their prey.

## Introduction

*Paryphanta busbyi* (Mollusca: Gastropoda: Rhytididae: Paryphantinae), commonly known as a kauri snail or pupurangi in Maori, is a large carnivorous land snail found in the northern part of the North Island of New Zealand (Fig. 1). Due to restricted distribution, nocturnal habits, protected conservation status and rarity, studies on the snail are infrequent. A one paragraph description of a preserved specimen was done by Hutton (1881) and in more detail by Murdoch (1903). Further studies established a phylogeny partly related to their distribution with a species recovery plan in mind (Powell 1946, Parrish 1992, Parrish et al. 1995). Spencer et al (2006) resolved phylogeny using mDNA COI sequences and despite latent morphological debate around two recognised subspecies *P. busbyi watti* and *P. busbyi busbyi* we accept Spencer’s phylogeny which does not distinguish them. Harmonic radar tracking of the snails developed by Lovei et al. 1997 helped answer some of the basic questions about the snail’s movements and biology with studies by Stringer & Montefiore (2000), Stringer et al. (2002), and Stringer et al. (2003).

**Fig. 1.**
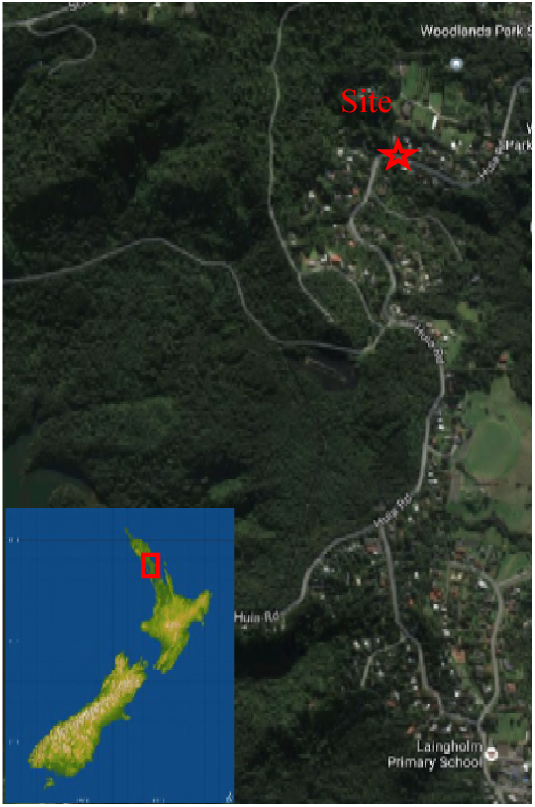
Map of New Zealand showing the location of the study between park and low density suburbia.

Studies of movements of *P. busbyi* using harmonic radar to relocate individual snails required recapture to confirm their presence and identities (Stringer & Montefiore 2000; Stringer et al. 2003). From the 126 snails included in Stringer et al.’s 2003 study insights into life history such as reproduction and mortality were gained. Their observations show the description of behaviours averaged over 126 individuals. The current study involves daily observations on a smaller number of snails, attempting to fill some behavioural gaps left in previous studies by long intervals between observations. Frequent observations meant minimizing the disturbance of the snails and their habitat had to be a priority for this study. It was hypothesised this approach would allow information to be gathered on *P. busbyi* that would elucidate some new aspects of the snail’s behaviour and put those already discovered into a better context.

## Materials and Methods

### Study site

The population of snails studied resided in the Woodlands Park, Waitakere ranges, Auckland, North Island, New Zealand (36° 57’ 42.82′S, 174° 37’ 10.38′E) (Fig.1).

The area was originally covered in temperate rainforest with mixed podocarp species, but was logged of nearly all large trees about 90 years or more prior to the study, leaving a few remnants of larger decaying unidentifiable stumps and log pieces. Attempts to convert the areas to farmland further altered the vegetation, which has now been regenerating for about 62 years. Grazing by brushtail possums (*Trichosurus vulpecula*) has resulted in the regeneration to a dominant canopy of kanuka (*Kunzea ericoides)*. While some larger pururi *(Vitex lucens)*, totara *(Podocarpus totara)*, rimu *(Dacrydium cupressinum)* and other species were left by loggers, most are currently small in stature. Temperatures in the area range from 6°C on a cold winter night to 22.5 or in microenvironments 28°C on a hot summer day. Frosts only occur in winter in localised exposed areas of ground, not among the trees. Rainfall is highly variable depending on local geography with no localized rain gauge. The nearest monitored rain gauge is just under 2000m away indicates rainfall is in the order of 2000 mm/yr in the area the snails were located.

More particular to the study area we have surroundings that form part of the eastern side of a north-south ridge. Snails were located and observed when they appeared in a shallow trench next to or more rarely on a 33.5 meters long stretch of exposed aggregate driveway. The trench is approximately 300mm wide, between 0 to 220mm deep originally for draining a small bush clad mainly south facing 30m long slope along 23m of the driveway’s length. The drain did not show an observable depth of water along its length during the year-long study. The only water observed was trickle fed from overhanging vegetation though during prolonged downpours it is likely to have more. On rare occasions snails were observed on much larger area of the driveway itself or on the opposite side of the driveway which has no formed channel and drops away down the hillside. Tunnels appeared periodically under the driveway.

### Daily snail observations

In order to know how often to observe the snail’s behaviour an indication of their speed of movement in the wild had to be determined. Speed in the context of snails in this paper is the time it took for a snail to cover a known distance without knowing if the snail moved continuously or intermittently. Even when the snail is videoed moving apparently continuously it is possible the snail’s mechanism of movement involved stopping and starting an unperceived number of times. The methodology used is not intended to elucidate the mechanism of movement but rather the overall distance covered over a period of time during different activities. Intermittent starting and stopping was easily perceptible during behaviours such as hunting or sensing the environment and may be more intuitively thought of as the snail’s pace for undertaking particular activities. The pace of most snails was established by revisiting snails over periods of minutes or sometimes hours and measuring their change in position using a measuring tape to determine how far the centre of the snail’s whorl had moved. For example, in the case of a 55mm snail we took a video on a hand-held Nokia RM978 phone with default automatic settings to determine speed. To determine the snail’s speed in the absence of the observer two still photographs were taken at a defined interval and the distance moved measured.

Searching for snails was done at least once a day during darkness using a red head light and occasionally during the day when the snails very rarely active. A total of 374 of the 377 days had at least one inspection. There is an unquantified bias towards observing larger snails as these are much easier to see and find. In at least 72 cases follow up inspections of discovered snails was done the same night to determine behaviours. Red light was used during night time photography. Measurements were done using a measuring tape without making contact with the snail. Avoiding contact of the measuring tape with the snail meant that most snail size measurements were within 5mm. Snails that were moving in a single direction would have their speed recorded by taking repeated measurements over minutes to hours depending on their speed, terrain and the ability to find the snail again without disturbing their habitat. For the purpose of this study snails were distinguished by their differing shell sizes and relative locations. Finding two snails of the same size on the same night only occurred once during the entire study and this pair were within 100mm of each other and small enough to be freshly hatched from the same nest. Smaller snails were more easily lost than larger ones as displacing the leaf litter in which they moved to relocate them was considered disturbing the environment. A mirror or endoscope were necessary to see or photograph within most lairs. Sometimes more than a few seconds of exposure to red or white light could cause snails to begin retracting into their shells though this behaviour varied in a similar way to that described for earthworm prey retracting into burrows (Darwin 1881). There were some longer duration observations (30min) but these were generally avoided as they might alter the wild snail’s natural behaviour more than is necessary for the study. Most observations of snails lasted only long enough to establish where snails were, what they were doing and to allow taking of the occasional photographs.

## Results

### Determining a snail’s pace

The most documented example of determining an adult snail’s speed was a rare daylight sighting on the 21^st^ of July 2015 at 8:08 am. A 60mm snail was filmed using a hand-held Nokia RM978 phone with default automatic settings. The video lasting 1min 11sec may be viewed at: https://plus.google.com/u/0/photos/116507430608150673768/albums/6181306159640899073/6181306164563221042. The terrain being transited was a sloped artificial concrete surface of exposed red aggregate. At the time of filming abrupt breaks in the cloud brought increased light levels during an otherwise dark dawn. The snail moved towards an area containing lairs and leaf litter many centimetres deep. Approaching the snail more closely than about a meter led to increased movement of eye stalks and tentacles including their partial retraction, a similar response to that seen at times to snails illuminated with red LED light during night time filming. Filming positions of the camera which gave this response were avoided where possible.

Measurements of the shift of the centre of the snail spiral during the period of the video taken at 8:08am on the 21st July 2015 when the snail moved approximately 0.02 meters was consistent with a snail’ s pace of between 0.9 to 1.1 meters per hour. Two still photos taken at 8:31 am and 8:40 am showed the centre of the spiral shell moved 0.27 meters, giving an approximate speed of 1.9 meters per hour. There was no observable slime trail. Another snail of a similar size was measured at night travelling near the same place as the former snail a year later at a speed of 2.9m/hour. These are considered travelling speeds on a relatively regular substrate with lesser speeds in leaf litter noted at other times later in this paper.

### A Selected short term observation

To help the reader interpret snail behaviours indicated in the year-long study a short period of the study is considered first in some detail. In fig. 2 a 42mm snail was first observed on the night of the 6^th^ of October 2015 and moved as indicated on the y axis. Circumstances meant that this 42mm snail was active and able to be tracked on more consecutive nights than any other snail. The movements seen in fig. 2 are consistent with the types of movements seen by other snails tracked over shorter periods during the rest of the year.

**Fig. 2.**
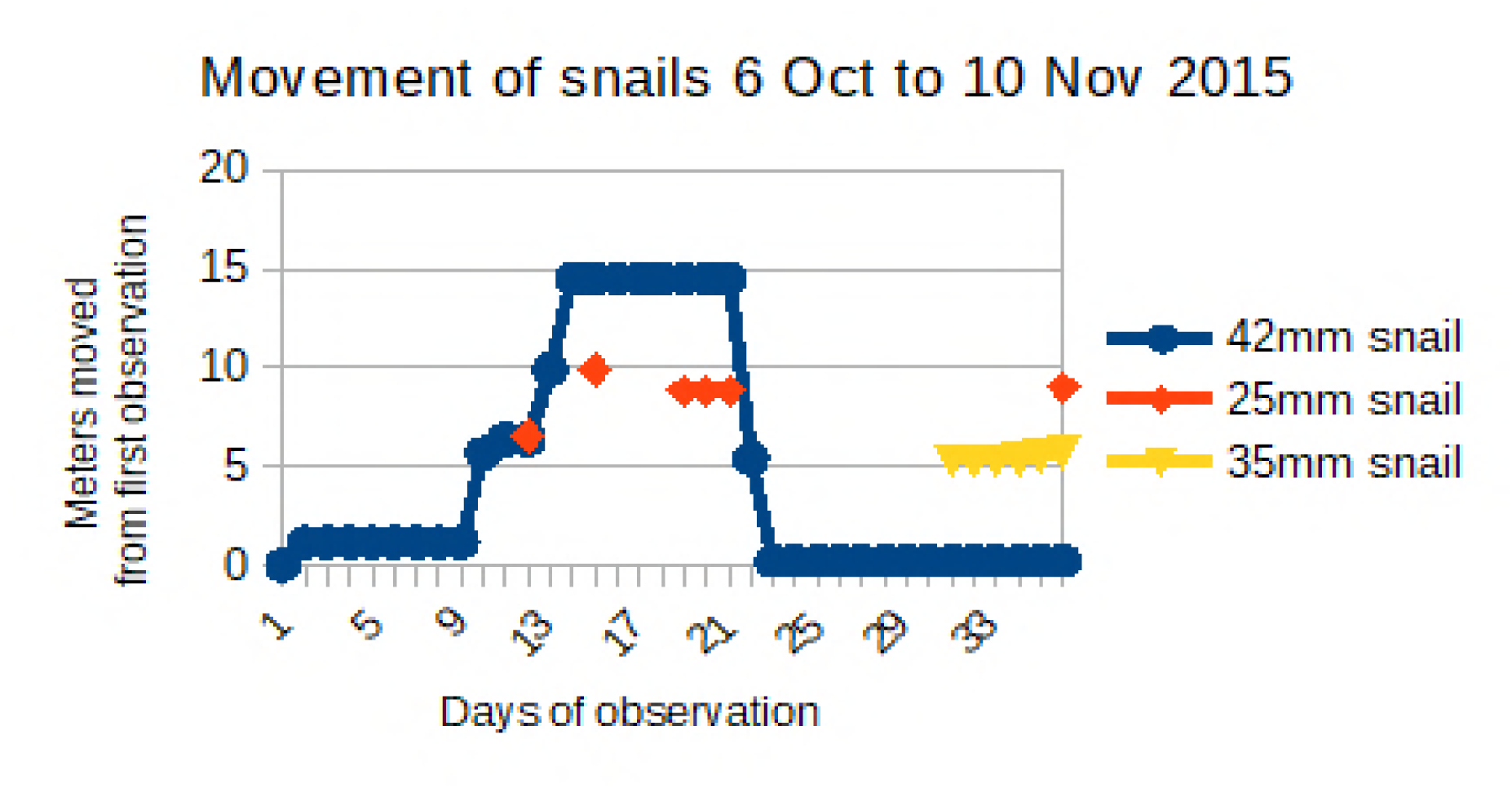
The distance moved in meters 3 snails from 6 October to 28^*th*^ October 2015 with the data accrued from 179 inspections over the 45 days.

During night time observation on day 1 the 42mm snail would appear to be hunting in deep leaf litter. It would move at rates of 0.5 to 1.5m per hour. While hunting the snail would wave its eye stalks and tentacles, pausing frequently and sometimes thrusting its head down into the deep leaf litter in any one spot for more than 10 minutes at a time (Fig. 3).

**Fig. 3.**
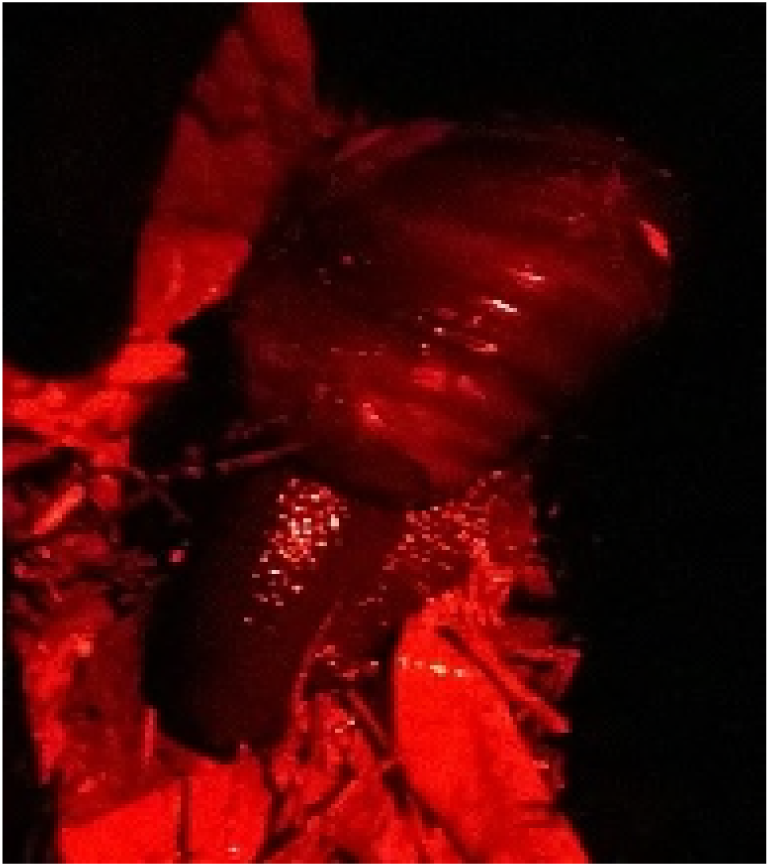
Snail photographed under red light from above. The head and neck are below the leaf litter curving downward away from the viewer with the shell and foot remaining at the leaf litter surface.

In this mode the snail moved about 170mm over 2 hours from approximately 22:15 to 01:00. By dawn the snail had moved into a lair 1.14 m from its original position. As indicated in fig. 2 by the horizontal graph lines the snail was observed in the lair day and night for days at a time. Repeated observations within the lairs showed the body partially extended towards the entrance of the lair, with eye stalks and tentacles sometimes extended, other times with the head retracted into the extended body. During daylight of day one the snail was seen probing the floor of the lair with its tentacles and also reaching up to probe the ceiling in a similar manner. The snail sometimes moved further towards the back or far left of the lair making it difficult to observe. While other snails observed in lairs were at time retracted into their shells it was clear that this snail was not retracted into its shell at any of the dozen or more times it was seen in the first lair it entered, even when the snail was motionless for prolonged periods. To summarise the activity within the lair the snail was sometimes stationary facing the front of the lair, sometimes mobile and sometimes less observable towards the back or not visible at all.

The snail was observed intermittently near the entrance of the first lair until day 6. Daily and nightly searches and lack of disturbance of obstacles put at the lair’s entrance indicated the snail did not emerge from the lair until day 8 after significant rain. All indications were that the snail remained in the lair for nine days though we cannot completely rule out short, unobserved forays outside the lair. When the snail was finally found outside the lair on day 9 it had moved 5.5 meters from the lairs entrance in the 6 hours since the previous search, indicating a pace under 1 m/hr. Observing the snail for half an hour while it appeared to search for prey showed 0.3 meters of linear movement, indicating a pace while hunting of 0.6 m/hr.

Other snails shown on the graph would also at times raise their heads well above the substrate they were traversing, moving the head from side to side while waving their tentacles. They may also leave a lair for a few hours and then return the same night. The snails often appeared to be scanning the environment; sensing the air for food, moisture or some other airborne factor. Having paused to sense the environment they would then resume their travels or stay in the lair. In fig. 3 consecutive horizontal graph points usually indicates the snails being in a lair. Only a couple of snails in the year of study were observed in the same place on consecutive nights when outside a lair. The snails at times returned to a similar position within a lair after each night’s activity, often to within millimetres, indicating some ability to navigate.

Other than the commonly observed behaviours of sensing, travelling, hunting and residing in lairs there were more rarely observed behaviours interpreted here as burrowing (Fig 4 11^th^ Oct 2016), climbing branches (Fig 5 9 May 2016 10:10pm) and maintaining the shell exterior (Fig 6 12^th^ August 2016).

**Fig. 4.**
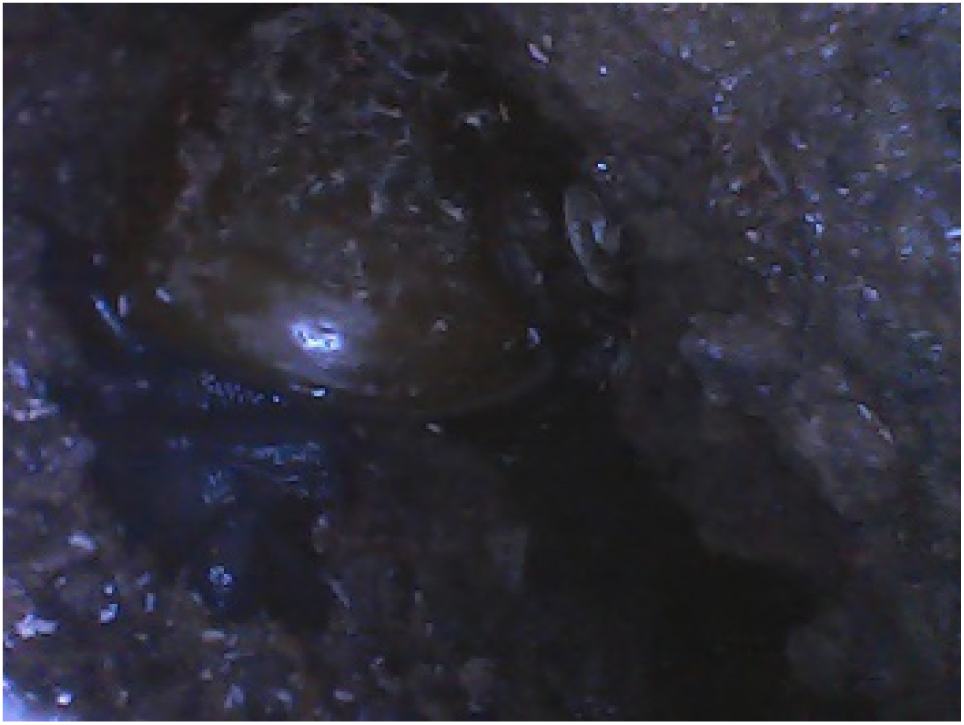
A snail burrowing photographed using an endoscope inserted into a lair. Mud is on the shell from where soil had contacted it. The tail is facing the viewer and appears wrinkled, perhaps as part of the burrowing action as it tries for force its way deeper.

**Fig. 5.**
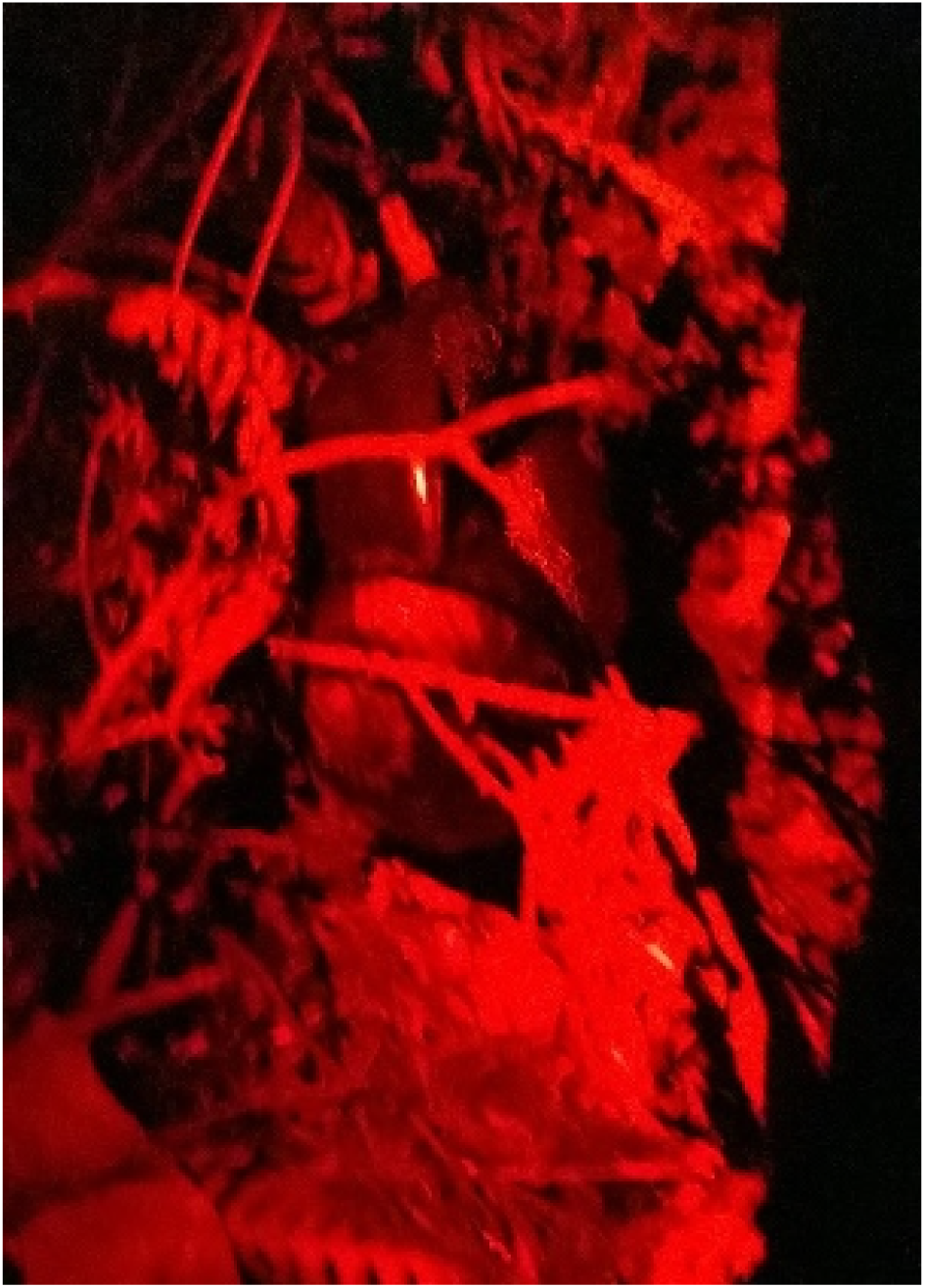
Snail climbing nearly vertically up a twig much narrower than the body of the animal. The back of the foot folded forward onto itself, wrapped around the twig. Snails climbing out of the study area on steep soil could fall backwards on lose soil. The stick apparently gives more reliable traction.

**Fig. 6.**
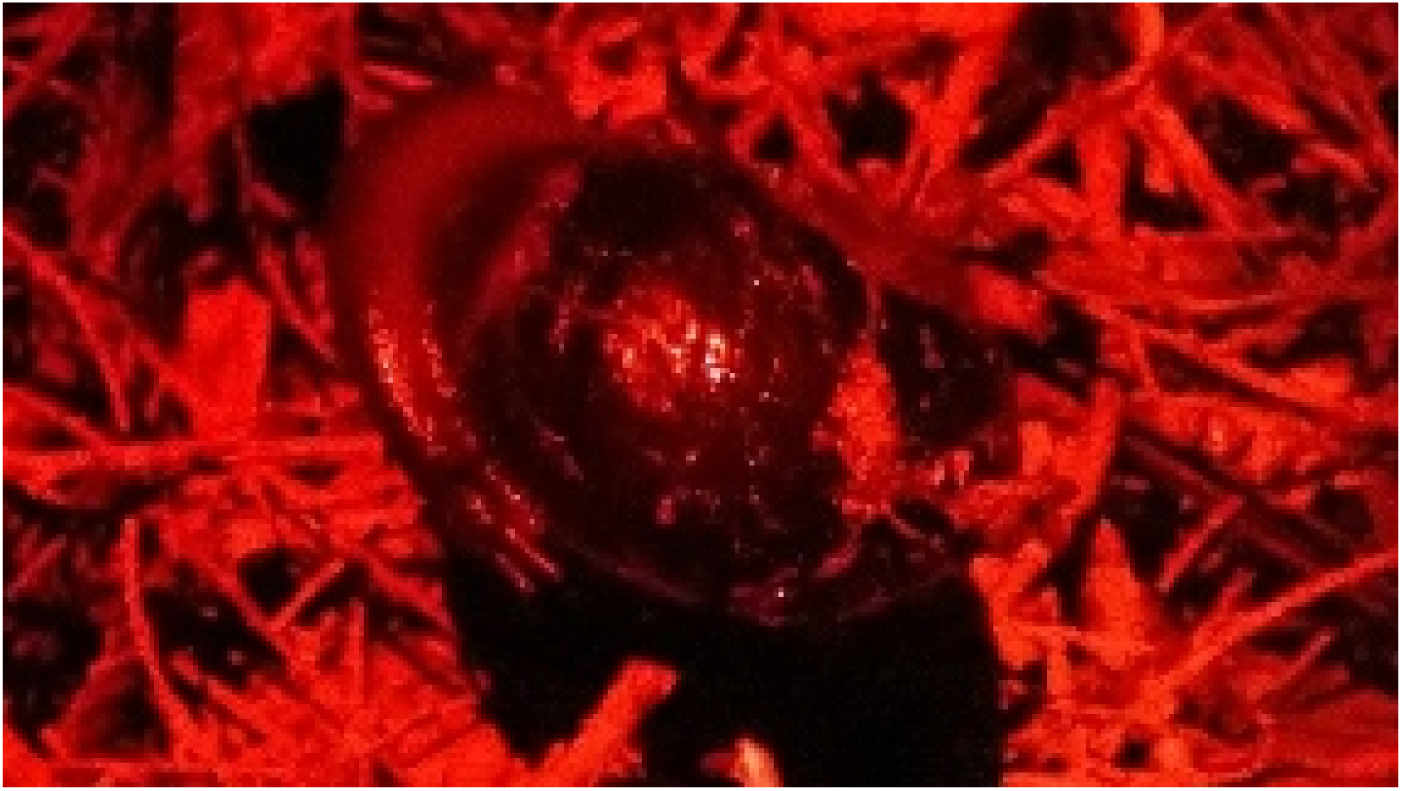
Snail with the head wrapped backwards over the shell. The shell appears to be covered in debris of some sort.

### Longer term daily observations

Data was collected for 372 days over a 377 day period with a total of 786 visits to the study area. The nine snails being plotted in fig. 7 is likely to under-represent the total number in the soil trench as the habitat was not destructively searched.

**Fig. 7.**
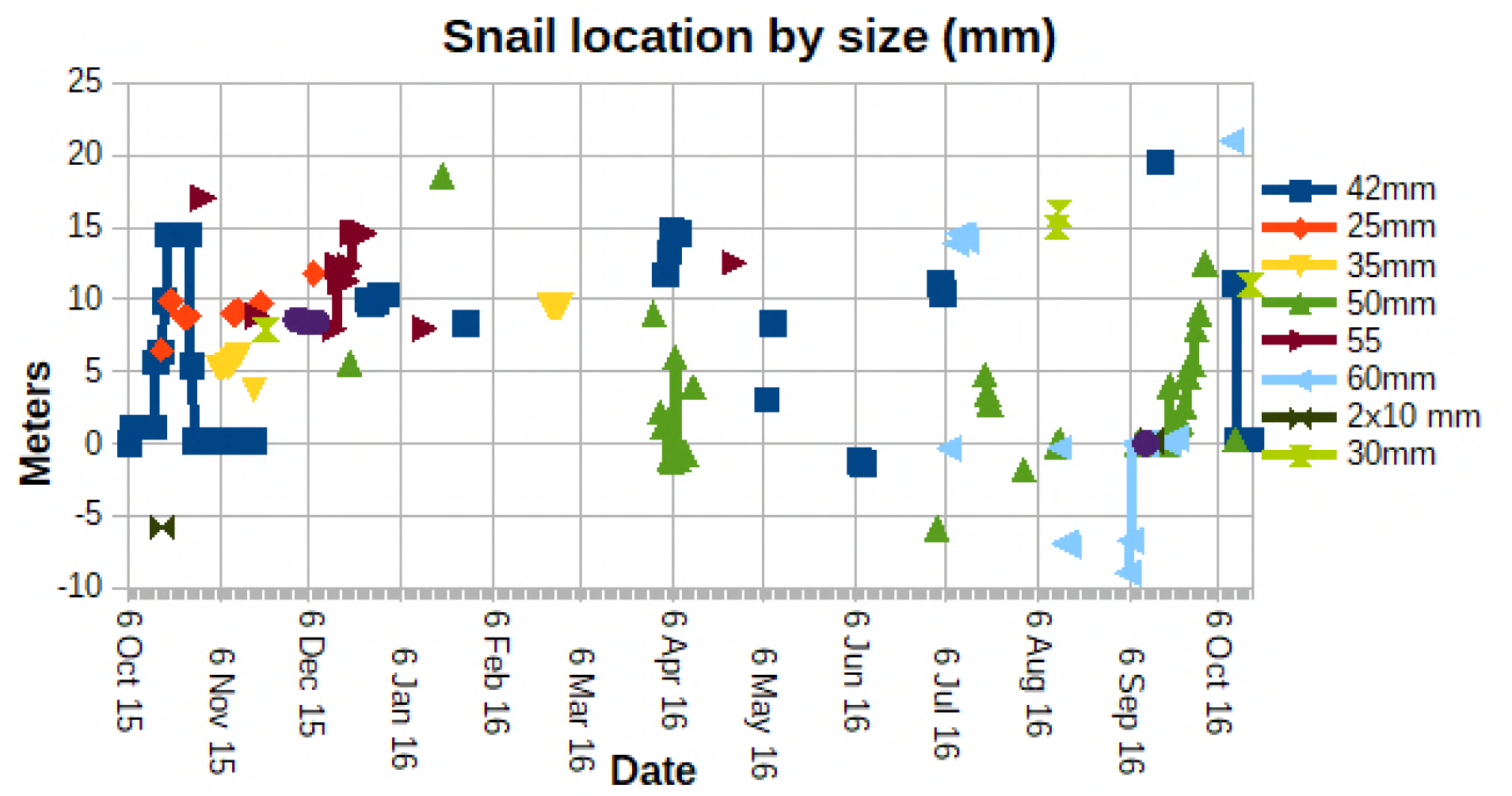
Just over a year of data showing the position of various sized snails in the study area at different times of the year. Where the lines connect the points the snail was observed frequently enough for the observer to ensure it is the same snail rather than just the same sized snail.

Over 372 days snails were observed on 191 days with 612 separate sightings recorded. Of the 191 days snails were observed they were in lairs on 100 days and outside lairs on 72 days. On 19 days they were observed at different times both inside and outside lairs or somewhere in between. The most common behaviours of no movement, sensing the air, travelling and hunting were difficult to define and are not quantified.

The data used to produce fig. 7 was elucidated further using Fisher Exact (P) tests suitable for rarer events as the snail number appearing on any one day were often 0 or 1. Comparing snail presence against local weather station data 1.73km away revealed significant correlations. The most significant factors were humidity (negative correlation P=0.000004359) and south west wind direction (150 to 200 degrees P=0.003784). There is an easily observable positive correlation in fig. 8 indicating numbers of snails being higher from the beginning of spring to the summer solstice and backed up by Fisher exact value P<0.0000001 comparing that period with the year as a whole. In summer and autumn the situation reverses (negative correlation P=0.00000369). Rainfall (P=0.4114) showed no significant correlation, with temperature (P=0.3113) and wind speed (P=0.2280) showing little more. Winter also shows no significant correlation with snail presence (negative correlation P=0.2692).

**Fig. 8.**
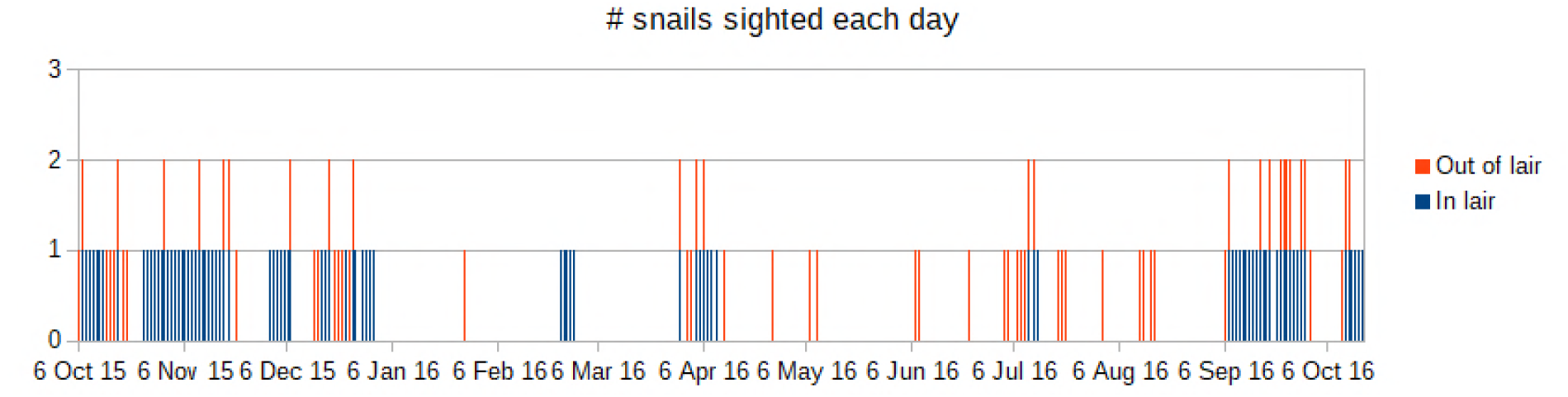
The number of observed snails in the study area was patchy over the course of the year, with numbers dropping to zero for weeks at a time over the driest and to a lesser degree wetter months. While there is apparently a small peak of activity near the beginning of each season the greatest activity is from early spring to the summer solstice.

## Discussion

Observations of wild *P. busbyi* for prolonged periods without disturbing the snails or the environment proved difficult. It is likely that snails in the study area were simply hidden too far underground or in the leaf litter to be observed. Despite this limitation it is reasonable to believe that at some time during the year the snails in the study area spend significant time underground day and night (fig.8). Lairs are at times used as a base with the snail leaving and returning to the same lair during a single night. At other times snails leave one lair for another. The significance of this newly observed behaviour may be speculated upon with possibilities including avoiding predators, avoiding light, or seeking areas of higher humidity. Underground environments chosen by the snails may attract suitable prey items and so allow daytime ambush of prey while the snails remain concealed. The ability to move between lairs is limited by the snail’s pace. If lairs are too far apart the snail may have to stay above ground during the day before reaching another lair. A maximum observed speed during this particular study of 2.9m per hour is more than the 2m per hour recorded in captivity for a similar carnivorous snail *Powelliphanta traversi traversi* (Devine 1997). There is no observable slime trail in *P. busbyi*. Slime trails are found in other snails which have been recorded moving at greater speeds e.g. garden snail at 3.5 to 5.4 meters per hour (Pavlova 2001, Mckee et al. 2013). In areas with suitable lairs greater than around 20m meters apart snails would be forced to spend time above ground during the day even if lairs are preferred. The ease with which smaller snails could find the smaller lairs may explain why they are difficult to relocate after being observed at the surface. Lair utilisation and burrowing would also explain the very low success rate in re-locating snails with harmonic radar as the radar does not penetrate the ground well (Lovei et al 1997, Stringer et al 2003). Any protection from predators snails might gain from using lairs may influence how large snails could grow as larger predators would more easily access the larger lair entrances larger snails would require to accommodate their shells.

Other behaviours of the snails observed included climbing branches much smaller in diameter than the animal itself while ascending steeper inclines, burrowing and the snail apparently maintaining the outside of its shell when it became dirty. While these activities may be expected and perhaps already observed by others they are documented via photographs here for *Paraphanta busbyi*.

Statistical analysis correlating snail activity with weather over the study period revealed the greatest significance of humidity followed by wind direction. The simplest hypothesis consistent with the humidity data is that the snails are present and most active in the study area when the humidity is lowest leading to the idea that the snails may prefer dry conditions. Such a hypothesis is easily refuted by the nature of the study area where the snails’ are preferring a trench with lairs on the south side of a hill where it is wetter than the surroundings the trench is acting as a drain for. The more likely valid hypothesis we will work with is that the drain is a humid, sheltered area preferred by the snails during periods of lower humidity outside the study area. A similar preference for humid conditions has been shown by another carnivorous snail (Roda et al 2016). The second slightly lesser correlation with wind direction is also consistent with the later hypothesis as on days the wind comes from the southeast the study area is not sheltered from the wind by the hill and no correlation with snail presence was found (Fisher Exact P=0.3502). South easterly winds funnelled up the valley will lead to greater agitation of the air in the study area causing the usually still and damp air trapped by the vegetation to become less humid. South-westerly winds in contrast seemed to correlate with snail presence (Fisher Exact P=0.003784) and this was also the direction of the main ridge in the area which acts as a barrier to the wind. The preference snails are showing for windless and humid conditions may be attributable to a generalization that molluscs prefer such environments. Snails can be used as indicators of soil humidity Čejka et al (2009) and different coloured snails of the same species may even respond differently to the same humidity Abdel-Rehim (2008). The ability for carnivorous snails to track prey more successfully in humid and windless conditions may play a role in this study. Having done the uncontrolled experiment of dropping an earthworm near the front of a snail and watching the reaction it is clear that, for minutes after the worm had “got away” straight past the snail and into a crevice the snail did a slow U turn and followed the path of the worm at a fraction of the worm’s speed. With the worm having disappeared down a crevice many minutes ago the snail eventually reached the crevice and forced itself as far as its shell would allow into the narrow gap the earthworm had entered. The snail stayed there for at least an hour, presumably waiting for the worm’s return. This demonstrable ability to track prey requires a systematic study of its own but would help explain why windless and humid air is preferred by the snails as such conditions would aid their ability to track prey and occupy good ambush positions. When trying to locate earthworms it may also be that the snail is using the strategy of finding areas where the preferred food of the earthworms is present. Earthworms themselves use smell to locate their food (Zirbes et al 2011) so it would be reasonable to hypothesis their predator does the same when trying to locate them. Another reason for the snails to visit the site is the requirement for calcium for the shell (Sulikowska-Drozd et al 2007). The driveway adjacent to the drain may be appealing to the snails as the aggregate used for concrete contained a few shell fragments as it is mined from a small island. No attempt was made to quantify this effect in the current study.

According to Lovei et al (1997) a *P. busbyi* snails ranged over about 1000 m^2^ during 2.5 years with the intervals of relocation varying from 6 to 16 weeks and displacements up to 25m from the most frequented area. It is unclear in the paper why the interval between relocations was so large and why only 1 of the 37 snails was relocated with that frequency. Similar limitations occurred in a much larger study by Stringer et al (2003) with 126 snails found and 93 fitted with transponders but the median recapture was 3 times over 26 field trips. They began their discussion “Many of our data are preliminary because we found and followed few of these rare snails and because of the long periods between observations”. From the current study it is worthwhile considering why the radar studies were problematic for recapture of the snails. This study shows the snails often frequent damp, humid, underground environments. Lovie et al (1997) noted “Water (also humidity) attenuated the signal.” of the radar with ground penetration of around 100mm. Snails more than 100mm underground may be hidden from the radar. In the current study snails were located with an endoscope up to 300mm into lairs and a large live snail was dug up 500mm below ground level in soil with high organic content in another site nearby (Kevin Barker, personal communication). With a measured snail’s pace in the current study of over 2m/hr and winter nights of over 12 hours 24m could be covered by a snail in one night. The long period between observations of months in other studies is therefore unlikely to give a reasonable representation of the snails range as there is ample time for snails to leave the area detectable by the radar and return again. From the data collected (Fig. 7) it appears the study site is being used on consecutive nights and by multiple snails per night as the bush starts to dry up at the beginning of spring until the summer sets in at the solstice. The snail’s preference for windless humid environments is clearly supported by the data and the study area is preferred at least in part for those reasons. The importance of humidity is hinted at by Stringer and Montefiore (2000) by showing a significant correlation between snail numbers and altitude. While not directly showing a correlation with humidity they noted cloud cover associated with the hill tops. We may postulate that snails not locatable in a study area may bury themselves deeper underground or they may leave the study area all together or they may do both. While not statistically defined snails have been observed leaving our study area. One was tracked for over 10m heading in a straight line away from the study site. Another was tracked a similar distance heading into the site. Fig. 5 was taken of a snail as it attempted to climb north out of the study site trench. During longer periods of no snail sightings an endoscope was used to search as deep into the lairs as possible most often without success in the mostly shallow lairs. It is reasonable to conclude the snails are moving in and out of the area as it becomes more or less appealing relative to the surroundings. This would make sense in terms of following their earthworm prey which are also known to migrate (Mather & Christensen 1992).

Lair based activity in windless humid environments has important implications for conservation strategies in the forest types studied here (Walker 2003). Ideal areas would require not only prey but year round high humidity, shade, minimal wind, the opportunity to burrow and good lairs to establish a thriving population. Such repopulation of suitable bush areas is unlikely to happen from what we know of their dispersal. It may well be that extinct birds such as the many varieties of Moa aided the snail’s dispersal to fragmented favourable areas in the past in a similar way that birds currently distribute other snails (Kazuto et al. 2008; Kramarenko 2014; Wada et al. 2014). Large Moa may have eaten snails whole with some passing through the digestive tract alive and so be deposited into new areas in their faeces. People can also act as good agents for dispersal (Gerber and Clark 2015). To prevent extinction of *P. busbyi* (IUCN red list: lower risk/near threatened) and other endangered snails qualified people may need to take an active role in assisting snails to colonize suitable new areas at speeds greater than a snail’s pace.

## Conclusion

*Paryphtanta busbyi* appear to be a very active predator moving at up to 2.9m/hr when travelling. Where lairs are present they may be used continuously for days at a time. When conditions are less suitable elsewhere the snails migrate to areas of a higher humidity and little wind, conditions that favour tracking their prey by smell. Such migrations lead to localized peaks in snail numbers and activity. With discontinuous studies these migrations may have led to miscalculations around the size of territories and estimates of population numbers. The ability of adult snails to spread to new areas is limited by their speed, quite specific environmental needs for hunting and lack of a known natural dispersal mechanism.

